# Condition-dependent fitness effects of large synthetic chromosome amplifications

**DOI:** 10.1101/2023.06.08.544269

**Authors:** Abigail Keller, Lucy L. Gao, Daniela Witten, Maitreya J. Dunham

## Abstract

Whole-chromosome aneuploidy and large segmental amplifications can have devastating effects in multicellular organisms, from developmental disorders and miscarriage to cancer. Aneuploidy in single-celled organisms such as yeast also results in proliferative defects and reduced viability. Yet, paradoxically, CNVs are routinely observed in laboratory evolution experiments with microbes grown in stressful conditions. The defects associated with aneuploidy are often attributed to the imbalance of many differentially expressed genes on the affected chromosomes, with many genes each contributing incremental effects. An alternate hypothesis is that a small number of individual genes are large effect ‘drivers’ of these fitness changes when present in an altered copy number. To test these two views, we have employed a collection of strains bearing large chromosomal amplifications that we previously assayed in nutrient-limited chemostat competitions. In this study, we focus on conditions known to be poorly tolerated by aneuploid yeast—high temperature, treatment with the Hsp90 inhibitor radicicol, and growth in extended stationary phase. To identify potential genes with a large impact on fitness, we fit a piecewise constant model to fitness data across chromosome arms, filtering breakpoints in this model by magnitude to focus on regions with a large impact on fitness in each condition. While fitness generally decreased as the length of the amplification increased, we were able to identify 91 candidate regions that disproportionately impacted fitness when amplified. Consistent with our previous work with this strain collection, nearly all candidate regions were condition specific, with only five regions impacting fitness in multiple conditions.

## Introduction

Aneuploidy, the presence of an abnormal number of chromosomes in a cell, can have contradictory effects on cellular fitness depending on environmental context. In multicellular organisms, aneuploidy can result in serious medical consequences such as sterility, intellectual disability, neuropsychiatric disorders, and spontaneous abortion (Yurov et al. 2018; Rutkowski et al. 2017; Potapova and Gorbsky 2017). Whole chromosome aneuploidies are often lethal in vertebrates, with only the autosomal trisomies of chromosomes 13 (Patau Syndrome), 18 (Edwards Syndrome), and 21 (Down Syndrome) capable of survival in humans beyond the first weeks of infancy. But while aneuploidy can be devastating on an organismal level, these same chromosomal abnormalities may contribute to the highly proliferative nature of cancer cells. Aneuploidy, along with chromosome instability (CIN), is extensively seen in tumor cells, to the point of being considered a hallmark of cancer, occurring in nearly 90% of solid tumors and 75% of blood cancers (Weaver and Cleveland 2006; Mitelman Database of Chromosome Aberrations and Gene Fusions in Cancer, https://mitelmandatabase.isb-cgc.org/). The tumorigenic nature and proliferative success of certain cancers may result from aneuploidies that alter expression of tumor suppressor genes or oncogenes, though causal explanations for this link are still limited in number (Lengauer et al. 1998; Williams et al. 2008; Sotillo et al. 2007).

While it is still in debate whether aneuploidy is a cause or consequence of unchecked cell growth in cancer, recent studies have reported links between the presence of aneuploidy and poor prognosis in cancer patients. Aneuploidy and CIN are associated with a higher chance of lethality in prostate (Stopsack et al. 2019) and breast cancers (Oltmann et al. 2018), as well as invasive behavior in Drosophila models of epithelial cancers (Benhra et al. 2018). Colorectal carcinomas tend to have a characteristic chromosome 7 amplification which is associated with enrichment in signaling pathways crucial for malignant transformation (Braun et al. 2019). Despite the correlation between prognosis and aneuploidy, it remains unclear whether tumor malignancy is dependent on simply the state of aneuploidy or the specific genes that have an altered copy number.

Single-celled organisms and fungi in particular (reviewed in Gorkovskiy and Verstrepen 2021) also show a complex relationship between aneuploidy and fitness. Cells with large CNVs are known to exhibit a slower growth rate in rich media, yet CNVs regularly arise under strong selection in stress environments. Budding yeast bearing specific chromosome abnormalities routinely evolve and outcompete euploid ancestors in nutrient-limiting experiments (Dunham et al. 2002; Gresham et al. 2008), and at least one chromosomal rearrangement has been directly linked to resistance to the antifungal drug fluconazole in *Candida albicans* (Selmecki et al. 2006). *Candida* and *Leishmania* species, both well-known human pathogens, display a high degree of genomic plasticity that may influence drug resistance and infectivity (Selmecki et al. 2005; Laffitte et al. 2016), and systematically generated trisomies in *C. albicans* have a constellation of fitness costs and benefits (Yang et al. 2021). While more cases of CNVs linked to disease, drug resistance, or pathogenicity continue to be discovered, in very few instances do we completely understand the mechanism by which large or complex chromosomal aberrations confer a fitness benefit or detriment. The effects of specific aneuploidies have in some cases been pinpointed to causative genes, such as fluconazole resistant *C. albicans* driven by the amplification of *ERG11* and *TAC1* on chromosome 5L (Selmecki et al. 2008) and increased lysine uptake through the amplification of *LYP1* on chromosome 14 in a mutualistic *Saccharomyces cerevisiae* community (Hart et al. 2019), but the majority of cases in both single- and multi-celled organisms are complicated by the sheer number of genes present in an abnormal copy number and the interactions they may have with one another. For example, candidate gene approaches have been slow in finding causative loci for many CNV-disease associations, and studies are biased towards candidate genes with already known or hypothesized links to the disease of interest (Usher and McCarroll 2015).

In our prior work, we addressed this problem in a yeast model by engineering a set of synthetic amplifications tiled across the entire *S. cerevisiae* genome (Sunshine et al. 2015). The strains in this telomeric amplicon, or “Tamp” collection, all contain a single amplification starting at a barcoded gene knockout from the heterozygous yeast gene deletion collection and extending to the telomere of the same chromosome arm (Figure 1A). The tiled nature of the Tamps revealed a step-like pattern when plotting the fitness of each strain by the chromosomal coordinate of the amplification start site, indicating that the fitness effects of amplifications change very little until particular regions of the chromosome are included in the amplification. This result is consistent with the hypothesis that in a growth condition where specific large amplifications have been shown to be beneficial, fitness effects could be conferred by a single or small number of genes that have been amplified. This previous study revealed that adaptation to nutrient limited environments through chromosomal copy number alterations was, in fact, largely driven by a discrete number of driver genes.

**Figure 1:**
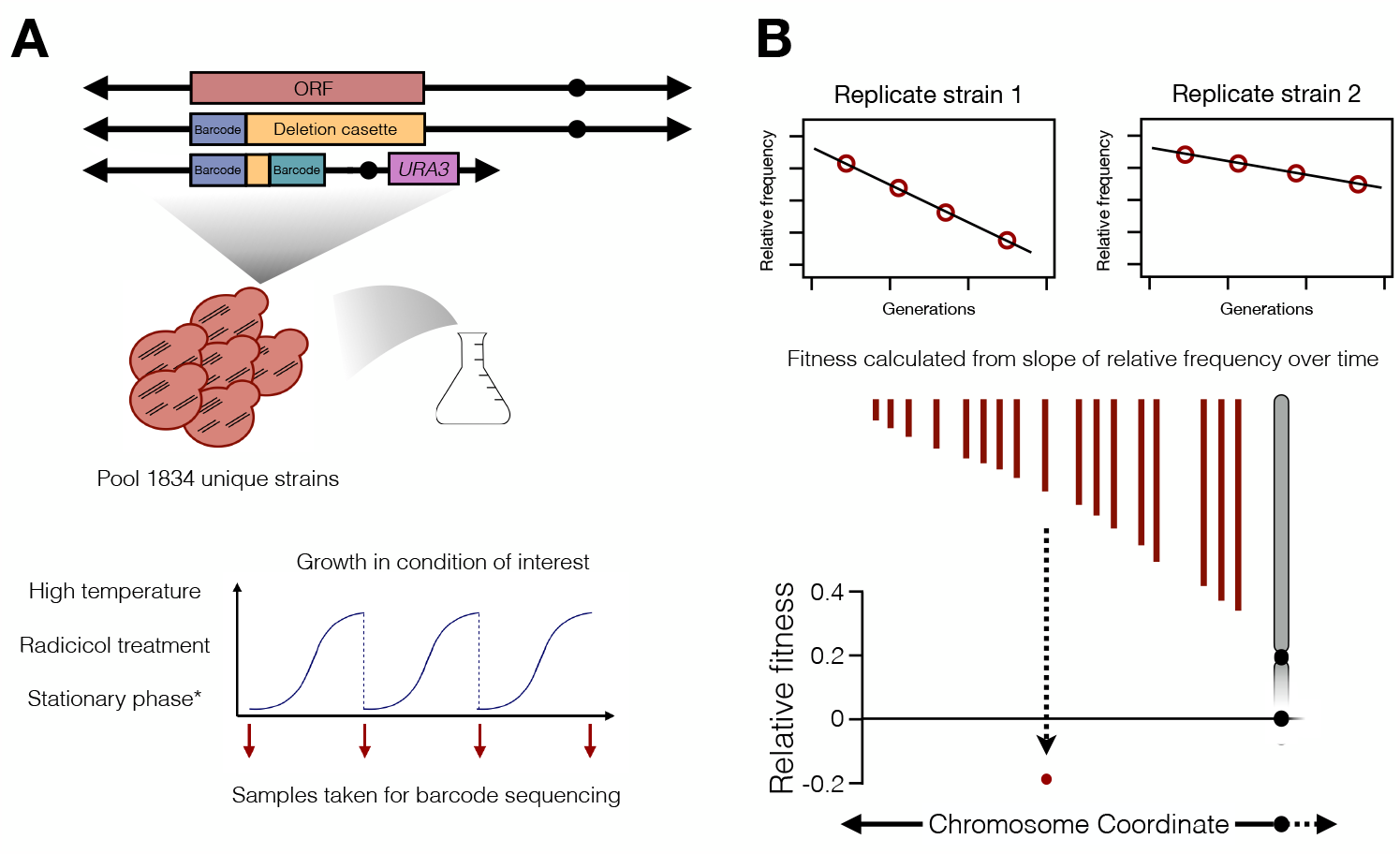
Overview of Tamp collection and competitions. (A) Overview of Tamp pool and competition experiments. Tamp strains were constructed by introducing a chromosome fragmentation vector bearing a region of homology to the deletion cassette in the yeast knockout strains. The resulting pool of Tamp strains were competed in batch culture. *The stationary phase experiment consisted of one growth curve and repeated sampling over 15 days while the culture remained in stationary phase (see methods). (B) Fitness for each Tamp replicate was calculated as the slope of the change in relative frequency over time. Final Tamp fitness is determined by the median fitness of all replicates for the strain.

While this result is in agreement with previous studies tracing the fitness benefit of amplifications back to specific genes in nutrient limited environments or in the presence of antifungal drugs (Payen et al. 2014; Gresham et al. 2008; Dunham et al. 2002; Linder et al. 2017; Selmecki et al 2008), the conditions tested are ones in which certain large amplifications are known to provide an adaptive benefit. In contrast, a survey of 1,011 natural *S. cerevisiae* isolates found a consistent fitness advantage of euploids over aneuploid and polyploid strains when tested across 36 growth conditions (Peter et al. 2018), though certain lineages of strains appear to be more tolerant than others (Scopel et al. 2021). Additionally, disomic yeast have been shown to share a set of common phenotypes such as increased glucose uptake and sensitivity to protein turnover and homeostasis perturbation due to the increased burden of expressing extra genes on large amplifications (Torres et al. 2007; Terhorst et al. 2020; Chunduri and Storchová 2019). The phenotypes exhibited by disomic strains were later shown to be accentuated by a mutation in *SSD1* in the *w303* strain background that disrupts proteostasis, highlighting the importance of both genetic background and proper protein homeostasis on the fitness of aneuploid cells (Hose et al. 2020).

Here we expand our Tamp screen to include conditions that aneuploid yeast tolerate poorly to investigate whether single genes disproportionately drive fitness effects in detrimental conditions, or if fitness deficits are largely due to the excess expression of many amplified genes. It is hypothesized that the general stress phenotypes exhibited by aneuploid cells results in a reduced fitness in high temperature, extended stationary phase, and treatment with Hsp90 inhibiting drugs (Torres et al. 2007). We found that larger amplifications are more detrimental to fitness in these three conditions; however, <100 discrete regions of the genome disproportionately drive these effects, showing that a complex combination of global and specific effects define the consequences of aneuploidy.

## Results

### Leveraging a Genome-Wide Collection of Telomeric Amplicon Strains

Amplification of a large chromosomal segment simultaneously changes the copy number and expression level of a large number of genes. In most cases, it is unclear how many of these genes contribute to the fitness effect of the amplification in a particular growth condition, or how this subset might change in a different environment. We have observed large amplifications rising to high frequency in laboratory evolution experiments under nutrient-limited environments, indicating that specific chromosome rearrangements and copy number changes may be beneficial to growth in these conditions (Gresham et al. 2008; Dunham et al. 2002; Kvitek and Sherlock 2011; Payen et al. 2014). Our previous work with the Tamp pool provided additional evidence that the effects of large amplifications in nutrient limitation are the result of a small number of genes, rather than the additive effect of many genes being amplified at once (Sunshine et al. 2015).

In this study we have expanded our focus to environmental conditions where aneuploidy is often detrimental to growth and survival: extended stationary phase, high temperature, and treatment with the Hsp90 inhibitor radicicol (Figure 1A). The mass overexpression of genes present on an amplified chromosome has been shown to result in stress on the cell’s protein homeostasis pathways by drastically increasing the pool of proteins that must be folded and later degraded (Torres et al. 2010; Pavelka et al. 2010; Dephoure et al. 2014). It has been hypothesized that this general stress is the basis for aneuploid cells’ increased sensitivity to conditions such as high temperature and drugs inhibiting Hsp90 that place additional strain on protein turnover pathways. Using the Tamp collection, we can see the degree to which these detrimental phenotypes are due to a small number of driver genes, as we predicted and observed in nutrient limitation, or due to the generalized overexpression of many genes at once.

We performed competitions of our Tamp pool in three conditions where yeast cells bearing whole chromosome amplifications have been shown to proliferate more slowly or lose viability faster than euploid yeast—high temperature, prolonged culture in stationary phase, and treatment with radicicol. These competition experiments were performed in batch culture, with multiple samples taken for barcode sequencing across approximately 17 generations of growth. Fitness measurements were calculated from the change in relative frequency of each strain across generations (Figure 1B, methods) derived from a linear slope. From these experiments, we gathered fitness data from ~1500 unique Tamp karyotypes across each condition with each karyotype represented, on average, by 18 independent replicates from the original construction of the pool (Supplemental Figure 1). The fitness for each Tamp strain was determined as the median value of these replicates in each condition (Supplementary Table 1). Because the Tamp collection was constructed from the heterozygous deletion collection, we performed an additional competition with this pool of deletion strains and excluded any Tamp strains where the corresponding deletion strain displayed a significant fitness effect (Supplementary Table 2).

### Chromosomal amplifications largely decrease strain fitness

We first looked at the general trend of fitness measurements as amplification size increased across each condition (Figure 2). A linear trend with a decreasing slope would indicate that fitness largely decreases as length of the amplification increases and that the specific genes present on amplifications matter less than the number of genes that have been amplified. Across most conditions, we found that this trend was approximately linear. On average, relative fitness decreased by -1.28×10^−4^ per kb across all conditions (−1.72×10^−4^, -3.22×10^−4^, 2.13×10^−4^, -1.88×10^−4^, and -1.72×10^−4^ for 30°C, 37°C, stationary phase, DMSO, and radicicol, respectively). Interestingly, prolonged stationary phase culture was the only condition that resulted in an overall positive correlation between fitness and length of amplification (Figure 2), while all other conditions resulted in a negative correlation. This is consistent with the hypothesis that large amplifications generally decrease fitness, unless condition-specific beneficial genes are present on the amplification.

**Figure 2:**
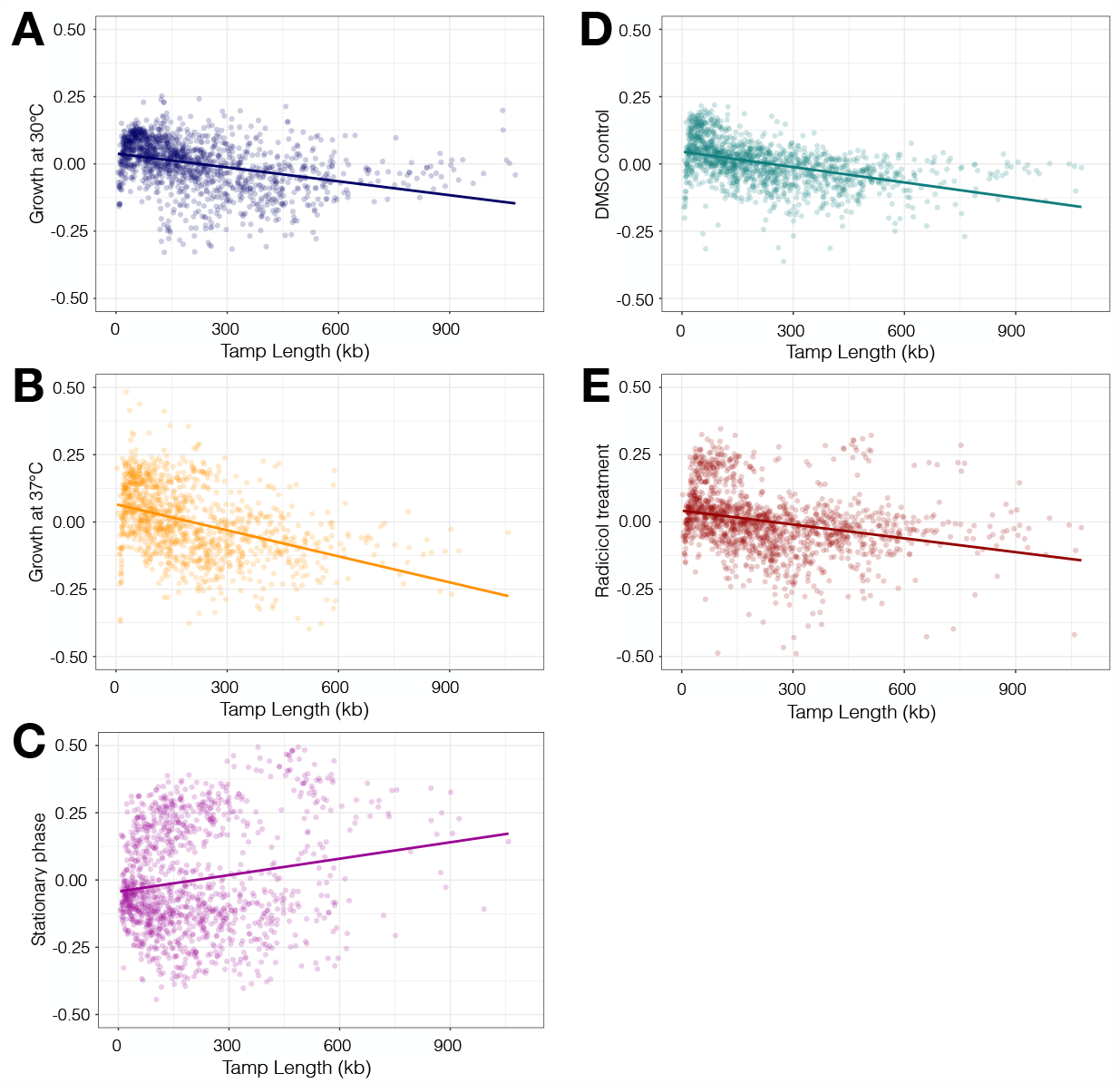
Fitness largely decreases as Tamp length increases: Fitness measurements of Tamp strains competed in (A) 30°C batch culture growth, slope = -1.72E-04; (B) 37°C batch culture growth, slope=-3.22E-04; (C) stationary phase, slope = 2.13E-04; (D) treatment with DMSO, slope = -1.88E-04; and (E) treatment with radicicol, slope =-1.72E-04.

We then separated out chromosomes to see if more complex patterns were apparent in different regions of the genome. Because all Tamps are designed to extend from a particular region in the genome to the telomere on the same arm, we also separated our fitness data by chromosome arm to more clearly observe potential patterns. Overall, 82.5% of chromosomes displayed a negative relationship between fitness and amplification length (15, 14, 8, 16, and 13 of 16 chromosomes for 30°C, 37°C, stationary phase, DMSO, and radicicol experiments, respectively). This general negative slope was also apparent when we separated the data on separate chromosome arms, as the amplifications in our collection should never cross the centromere. Across all conditions, 73.75% of chromosome arms displayed a negative correlation between amplification length and fitness (26, 25, 13, 24, and 30 of 32 chromosome arms for 30°C, 37°C, stationary phase, DMSO, and radicicol experiments, respectively).

While the overall trend remained that longer amplifications usually had lower fitnesses, multiple regions stood out as showing discontinuous patterns that deviated from the linear relationship between fitness and amplification length (Figure 3A, note locations where data in red do not follow the linear fit in black). Additionally, these regions did not behave similarly across different environmental conditions, indicating that amplifying particular regions of the genome has contrasting effects in these different cases (Figure 3B). To further examine this occurrence, we plotted each Tamp’s fitness by the chromosome coordinate of the amplification start site. In our previous work in nutrient limitation, this allowed us to observe a step-like pattern across chromosomes where fitness appeared to change very little until certain regions of the genome were amplified. We hypothesized that these regions contained genes that were disproportionately impactful on fitness when amplified in a particular environment. Visualizing our new dataset in this way is consistent with the hypothesis of small numbers of genes having a disproportionate effect on fitness that we previously observed in nutrient limitation.

**Figure 3:**
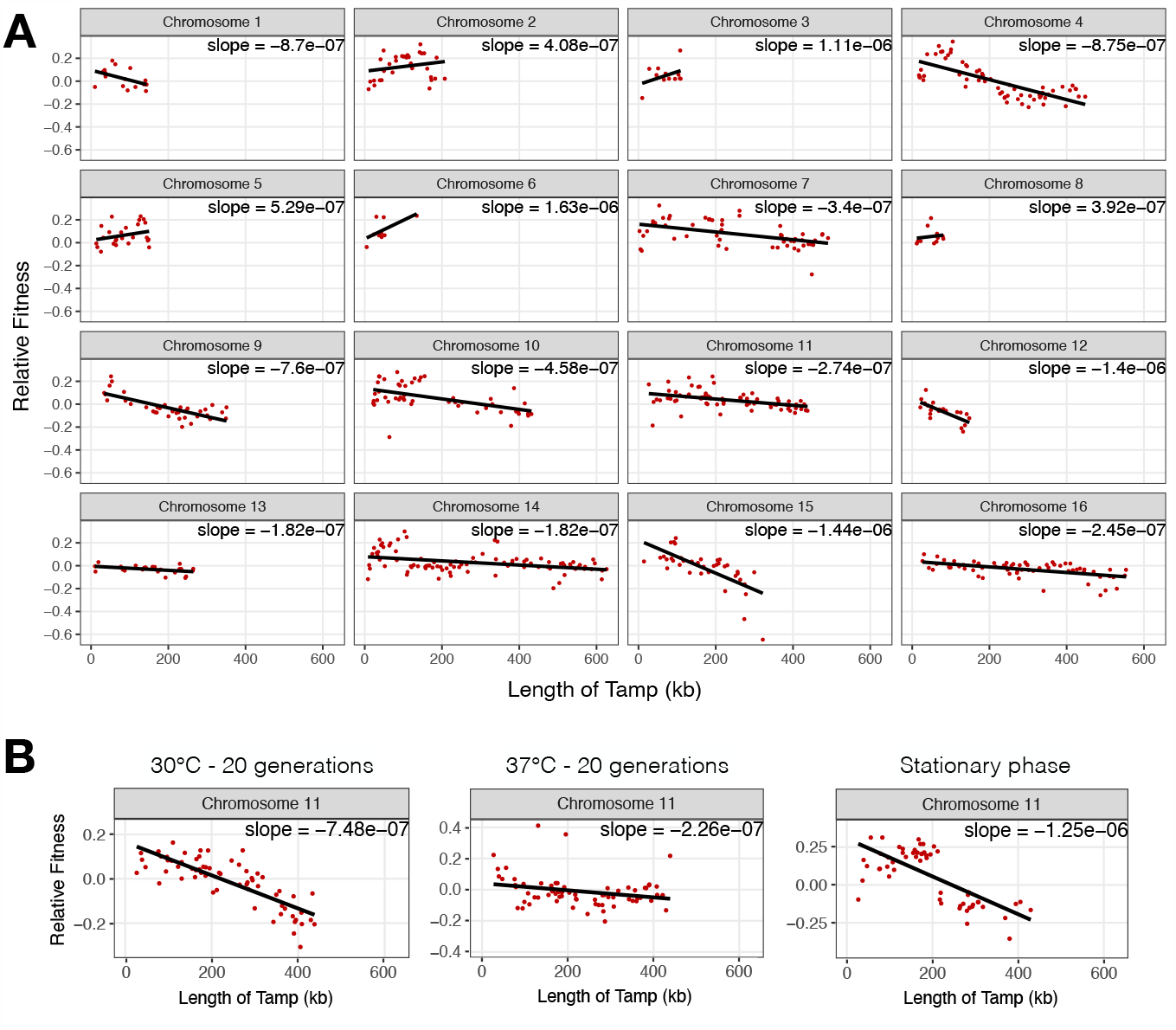
Nonlinear and condition specific trends in fitness across chromosomes: (A) Fitness of Tamp strains grown in 20 μM radicicol by length of amplification. Data are from the amplifications of the left arm of the chromosome only. (B) Fitness of Tamps with amplifications on the left arm of chromosome 11 from competitions in extended stationary phase and high temperature.

### Identification of genomic regions with high impact on fitness

To more quantitatively analyze the possibility of certain genes having a disproportionate effect on fitness in more detrimental conditions, we fit a piecewise constant model using the 1D fused lasso to our fitness data across each chromosome (Tibshirani et al. 2005). We selected the largest tuning parameter resulting in a piecewise constant model within two standard errors of the best fit. This is a relaxation of a more standard practice of selecting the largest within one standard error of the best fit. We employed this relaxation to limit overfitting of smaller step points resulting from experimental noise. This analysis allowed us to identify candidate “fitness breakpoints” where fitness of amplifications is disproportionately impacted by a small number of genes (Figure 4A). Using this method, we identified an average of 70 candidate step points that may contain genes of large effect for each experimental condition (Table 1). In all conditions except stationary phase, many more of the candidates were ‘downsteps’—regions where fitness decreased once they were present on the amplification.

**Figure 4:**
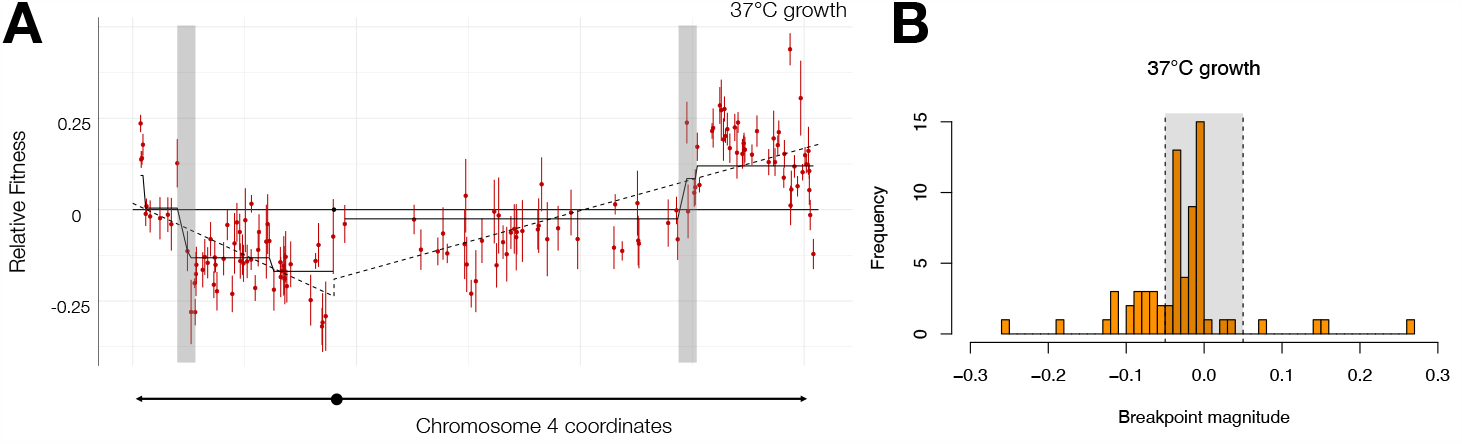
Piecewise constant model identifies candidate regions of large effect: (A) Fitness of Tamp strains with amplifications on chromosome 4 plotted by chromosome coordinate. Dotted line represents a linear regression model and solid line represents a piecewise constant model fit by 1D fused lasso. Shaded regions are breakpoints of high magnitude. (B) Histogram of breakpoints from competition in 37°C. Dotted lines represent cutoff values of +/- 0.05.

**Table 1.**
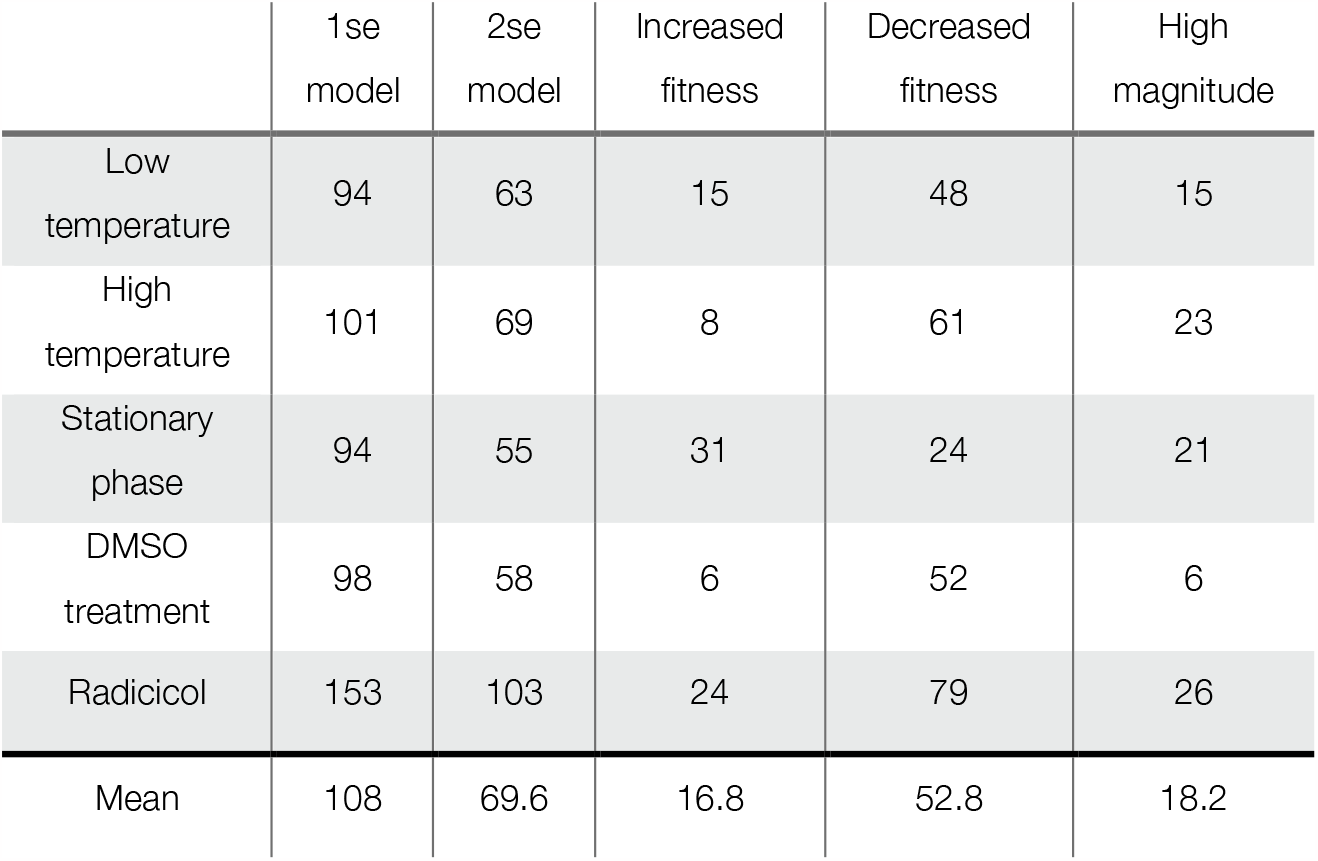
Summary of candidate break points from piecewise constant models.

|  | 1se<br>model | 2se<br>model | Increased<br>fitness | Decreased<br>fitness | High<br>magnitude |
| --- | --- | --- | --- | --- | --- |
| Low<br>temperature | 94 | 63 | 15 | 48 | 15 |
| High<br>temperature | 101 | 69 | 8 | 61 | 23 |
| Stationary<br>phase | 94 | 55 | 31 | 24 | 21 |
| DMSO<br>treatment | 98 | 58 | 6 | 52 | 6 |
| Radicicol | 153 | 103 | 24 | 79 | 26 |
| Mean | 108 | 69.6 | 16.8 | 52.8 | 18.2 |

While using a tuning parameter within two standard errors (rather than one standard error) of the best fit decreased the number of small steps present in the data from an average of 108 steps per condition to an average of 70, we still found a high number of smaller steps in the model, indicating we may still be overfitting the data and including steps that were most likely from experimental noise rather than those of biological relevance. In each condition, many of the steps had a very low magnitude with a much smaller number at higher magnitudes (Figure 4B, Supplemental Figure 2). To limit our initial pool of candidate steps, we chose to focus on the step points of highest magnitude. Selecting only steps with a magnitude of 0.05 or higher narrowed our candidate pool of step points to an average starting set of 18 regions per condition (15, 23, 21, 6, and 26 candidate regions for 30°C, 37°C, stationary phase, DMSO, and radicicol experiments, respectively, listed in Supplemental Table 3).

### Effects of aneuploidy in detrimental conditions are still largely condition specific

Our work investigating the effects of large amplifications in nutrient limited media suggested that the fitness benefits, or detriments, conferred by copy number increase of particular chromosome segments are largely condition specific. To test if this result holds true in our batch culture experiments, we compared candidate step points of high magnitude to see whether they were largely shared or unique to specific growth environments. Of 91 step points across all five conditions, only 5 regions were present in multiple conditions, and none were shared between all conditions (Figure 5). Furthermore, all but one of these shared candidate regions were downsteps, regions that decreased fitness when amplified. This finding suggests that while most positive and negative fitness effects of gene amplifications are still condition specific, detrimental phenotypes are more likely to be shared across multiple conditions. One exception to this observation is a region on chromosome 12 that resulted in an upstep in extended stationary phase and a downstep in high temperature. This region highlights an interesting case where amplification of a gene may have contradictory effects in these two conditions.

**Figure 5:**
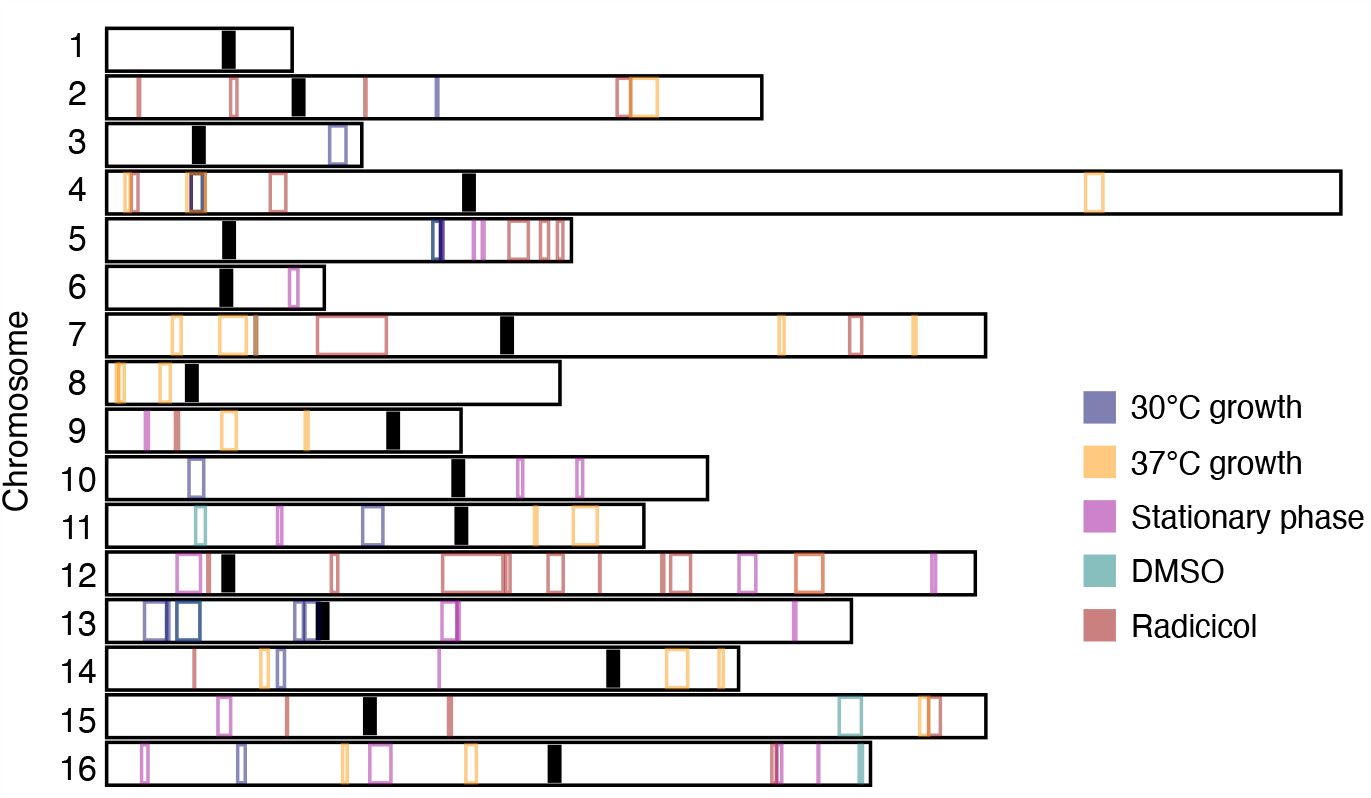
Regions of large effect identified by piecewise constant model are condition specific: Breakpoints with a magnitude greater than 0.05 as identified in piecewise constant model are outlined for each condition.

## Discussion

Large chromosomal amplifications are a relatively common error in mitosis and meiosis and can cause significant problems on a cellular and organismal level. However, studies in yeast species have repeatedly highlighted the complex and sometimes contradictory effects aneuploidy can have on fitness. While large amplifications are a frequent mechanism by which organisms can adapt to new environments, they also result in sensitivity to an array of environmental stressors. In yeast, amplifications of most whole chromosomes result in lowered tolerance of high temperatures, compounds impacting protein homeostasis, and prolonged stationary phase (Torres et al. 2007). It has been hypothesized that these environments exacerbate the stress on protein turnover pathways from expressing, folding, and degrading products of many amplified genes at once.

To investigate whether fitness deficits associated with large amplifications are dependent solely on the number of genes amplified or the specific genes present on the amplification, we used a collection of yeast with large amplifications tiled across the entire genome (Sunshine et al. 2015). Using this collection, we found that while fitness does decrease as length of the amplification increases, the correlation is weaker than would be expected if only the number of genes present on the amplification contributed to fitness. In some cases, fitness even increased with increasing Tamp length, suggesting the presence of one or more genes that provided an adaptive benefit when amplified in these conditions. By applying a piecewise constant model to our data, we were able to identify small regions of the genome that impact fitness in both positive and negative ways, indicating the negative effects of large amplifications may also be driven by individual genes in certain cases. This is consistent with our Tamp competitions in nutrient limitation, where we found discrete regions of the genome driving fitness effects.

It is important to note two potential limitations of data presented here - errors present in our Tamp pool and limitations of our model. As we have previously described, larger Tamps are more likely to have errors resulting from the pooled construction of the collection, including incomplete amplification extending all the way to the telomere. We have attempted to address this problem with the inclusion of multiple replicates of each Tamp karyotype and stringent filtering steps during analysis to exclude Tamps with high variability in fitness. However, it is possible that these errors may still be confounding our fitness measurements and models, particularly of Tamps on larger chromosome arms. The collection was also designed on the backbone of the yeast knockout collection, which has been shown to harbor mutations and chromosome abnormalities resulting from adaptation to gene deletions (Puddu et al. 2019). While this undoubtedly introduces noise into our experimental data, calling fitness breakpoints as the result of the pattern that many individual Tamp strains together form limits the impact of these errors somewhat. Second, despite allowing for a larger tuning parameter and simpler model to limit overfitting of our data, we still found that many fitness breakpoints called by the model were small and likely attributable to noise in the data. To account for this, we limited our focus of potential candidate genes to only those present in breakpoints of high magnitude. This leaves open the chance that we have filtered out true candidates whose effects were masked by experimental noise. The presence of many small breakpoints in the piecewise constant model could also be indicative of the likelihood that both length of amplification and gene content contribute to fitness. In this case, the many small breakpoints are trying to fit a general linear trend and are interspersed by larger breakpoints caused by large effect genes.

In this study we focus on the genes that may be impacting fitness when amplified, but other genomic features may also contribute to observed fitness effects. We have defined fitness here as a change in relative frequency over time, a metric that encompasses both the ability of cells to maintain viability and the relative speed at which they grow and divide. In order to divide, a cell must complete replication of all chromosomes. In yeast, this process initiates at autonomously replicating sequences (ARSs) and proceeds bidirectionally across chromosomes. Our Tamp collection relies on native ARSs to replicate our amplifications; however, long stretches of a chromosome without a strong or early firing origin of replication may impact the time it takes for a cell to divide. In this case, Tamp strains with amplifications extending into long stretches devoid of an early origin would show a decreased fitness compared to a strain with a slightly longer amplification encompassing an ARS, resulting in an upstep in fitness in our assay. The impact of replication origins and other genomic features such as transposable elements requires further investigation.

The contributions that individual genes make towards overall cellular fitness of aneuploid cells have implications even beyond how microbes may adapt to new environmental challenges. As discussed earlier, aneuploidy is exceedingly present in nearly all cancers. Additionally, cancers derived from different tissue types display different recurring amplifications or deletions of specific chromosomes (Kim et al. 2013). This suggests that specific genomic rearrangements may provide a proliferative advantage to cancers present in different parts of the body, possibly due to the amplification or deletion of specific genes impacting malignancy. This work lends strength to the hypothesis that negative impacts of aneuploidy may also be driven by copy number changes of individual genes, bolstering the possibility of drugs and therapeutics specifically targeted to the karyotype of the tumor. Because cancer cells have unique karyotypes often with many different chromosome copy number changes, they may be uniquely sensitive to specific compounds that target amplified or deleted genes in their genome.

## Materials and Methods

### Strains used in this study

Strains and oligos used in this study are listed in Supplemental Tables 4 and 5. The genome-wide Tamp pool construction is described in (Sunshine et al. 2015). Tamp strain competitions were carried out in minimal media +20mg/L histidine with additional additives if indicated. Heterozygous deletion strain competitions were carried out in minimal media with 20mg/L histidine and 20 mg/L uracil. Yeast were grown at 30°C unless otherwise noted.

### Plate reader experiments to determine radicicol concentration

The radicicol concentration used for Tamp competitions was determined by testing the effects of 0μM, 2.5μM. 5μM, 10μM, and 20μM radicicol on doubling time of wild type cells and cells bearing a large chromosomal amplification (AK2, AK3, AK4; Supplementary Table 4). Log-phase cells were diluted to an OD600 of 0.05 in appropriate synthetic media in a 96-well flat-bottom plate. Growth curves were obtained on a BioTek Synergy H1 Multi-Mode plate reader over the course of 48 hours. A concentration of 20μM radicicol resulted in greatest stratification of growth rate effects on Tamp strains with only modest effects on wild type cells (Supplementary Figure 3).

### Competition experiments

Competitions with the Tamp pool were performed in batch culture as follows. Competitions with corresponding strains from the yeast heterozygous deletion collection (Tong et al. 2001) were conducted in parallel in B+His+Ura medium using the same experimental procedures.

#### Temperature competitions (30°C and 37°C)

Two replicate flasks of B+His medium were inoculated with the Tamp pool for each temperature at an initial OD600 of 0.05. Cultures were grown in shaking incubators for approximately 17 generations, with back dilutions to an OD600 of 0.05 approximately every 36 hours. At each back dilution and at the conclusion of the experiment, 2 mL of culture were harvested and processed for genomic DNA using a modified Smash-and-Grab protocol (Hoffman and Winston 1987).

#### Stationary phase competitions

Two replicate flasks of B+His medium were inoculated with the Tamp pool at an initial OD600 of 0.05. Cultures were grown in a shaking incubator for approximately 2 weeks. Samples were taken for sequencing on days 1, 5, 9, 11, and 13. To avoid sequencing residual DNA from dead cells, 750 μL of culture were diluted in 4 mL of fresh medium and grown for approximately 16 hrs. 2 mL of culture were then harvested and processed for gDNA.

#### Radicicol competition

Two replicate flasks of B+His media with either 20 μM radicicol in DMSO or DMSO only (final concentration 2mM or 0.0015% DMSO) were inoculated with the Tamp pool at an initial OD600 of 0.05. Cultures were grown in a shaking incubator for approximately 19 generations, with back dilutions to an OD600 of 0.05 approximately every 31 hours. In addition to the 2 mL samples harvested at every back dilution and at the end of the experiment, samples of at least 2 OD units of cells were harvested at hours 10.5, 14, 17, and 23.5 and processed for gDNA.

### Fitness determination by Illumina barcode sequencing

Barseq libraries were prepared using primers OAK1-3 for the Tamp pools and OAK8-9 for the deletion pools (Supplementary Table 5) and purified using a Zymo Research DNA Clean & Concentrator kit (cat. no. D4004). Libraries were pooled at an equimolar ratio, size selected with Beckman Coulter Agencourt AMPure XP PCR Purification beads (item no. A63880) at a ratio of 1:1, and verified on Invitrogen Novex 6% TBE gels (cat. no. EC62655BOX), Agilent D1000 DNA ScreenTapes (cat. no.5067-5582), and/or a Roche KAPA Library Quantification Kit (kit code KK4828). Each pool was sequenced on an Illumina NextSeq, generating a total of 455,443,917 total reads across all conditions. Fitness calculations were made with custom python and R scripts (available at https://github.com/dunhamlab/Tamp-analysis) based on analysis approaches published previously and summarized below (Sunshine et al. 2015; Payen et al. 2016). Raw fastq files can be accessed at BioProject ID PRJNA725555.

Barcode reads were filtered using the FASTQ Quality Filter from the FASTX-Toolkit (http://hannonlab.cshl.edu/fastx_toolkit/) to eliminate barcodes containing any base with a Q score of less than 20. Remaining read pairs were assigned to a Tamp strain based on the 20-bp yeast deletion collection uptag or downtag, indicating genomic location of the *KanMX* cassette used as the Tamp initiation site, and an additional 12-bp barcode on the opposite read marking replicates of each strain. Deletion-replicate pairs were further filtered for sufficient number of reads with a cutoff of an average of 5 reads per time point, and if necessary, only the 100 replicates of each strain with the highest numbers of reads were used to determine fitness. Read counts were then normalized by taking the log_2_ ratio of the frequency of the replicate at each time point relative to the initial frequency in the pool. Read counts of zero were considered missing data. The log_2_ ratios were plotted against the generation time (or day number for the stationary phase experiment) yielding an average slope for each replicate. Fitness was determined as the median fitness among replicates. To account for possible incorrect karyotypes, such as truncation of the desired amplification, Tamp strains with high variability in replicate fitnesses, defined as having a standard error over 1 standard error above the mean standard error, were discarded.

### Identification of candidate driver genes

To identify chromosome regions of large fitness effect, we fit a piecewise constant model using a 1D fused lasso. Initially, we used 5-fold cross validation to select the largest tuning parameter that resulted in a piecewise constant model within one standard error of the best possible fit. While this method identified many large step points for further study, we found that in some cases, the model overfit the data and included step points not likely to actually include driver genes. To limit this overfitting, we relaxed the model, allowing for the largest tuning parameter within two standard errors of the best fit. We then prioritized step points of the highest magnitude for further investigation. Candidate step regions are listed in Supplementary Table 3.

## Acknowledgements

Thanks to Bonny Brewer for discussion of the impact of ARS density on Tamp fitness.

## References

Benhra N, Barrio L, Muzzopappa M, Milán M (2018) Chromosomal Instability Induces Cellular Invasion in Epithelial Tissues. Dev Cell 47:161–174.e4. https://doi.org/10.1016/j.devcel.2018.08.021

Braun R, Ronquist S, Wangsa D, et al (2019) Single Chromosome Aneuploidy Induces Genome-Wide Perturbation of Nuclear Organization and Gene Expression. Neoplasia 21:401–412. https://doi.org/10.1016/j.neo.2019.02.003

Chunduri NK, Storchová Z (2019) The diverse consequences of aneuploidy. Nat Cell Biol 21:54–62. https://doi.org/10.1038/s41556-018-0243-8

Dephoure N, Hwang S, O’Sullivan C, et al (2014) Quantitative proteomic analysis reveals posttranslational responses to aneuploidy in yeast. Elife 3:e03023. https://doi.org/10.7554/eLife.03023

Dunham MJ, Badrane H, Ferea T, et al (2002) Characteristic genome rearrangements in experimental evolution of Saccharomyces cerevisiae. Proc Natl Acad Sci U S A 99:16144–16149. https://doi.org/10.1073/pnas.242624799

Gorkovskiy A, Verstrepen KJ (2021) The Role of Structural Variation in Adaptation and Evolution of Yeast and Other Fungi. Genes (Basel) 12:699. https://doi.org/10.3390/genes12050699

Gresham D, Desai MM, Tucker CM, et al (2008) The repertoire and dynamics of evolutionary adaptations to controlled nutrient-limited environments in yeast. PLoS Genet 4:e1000303. https://doi.org/10.1371/journal.pgen.1000303

Hart SFM, Pineda JMB, Chen C-C, et al (2019) Disentangling strictly self-serving mutations from win-win mutations in a mutualistic microbial community. Elife 8:e44812. https://doi.org/10.7554/eLife.44812

Hoffman CS, Winston F (1987) A ten-minute DNA preparation from yeast efficiently releases autonomous plasmids for transformation of Escherichia coli. Gene 57:267–272. https://doi.org/10.1016/0378-1119(87)90131-4

Hose J, Escalante LE, Clowers KJ, et al (2020) The genetic basis of aneuploidy tolerance in wild yeast. Elife 9:e52063. https://doi.org/10.7554/eLife.52063

Kim T-M, Xi R, Luquette LJ, et al (2013) Functional genomic analysis of chromosomal aberrations in a compendium of 8000 cancer genomes. Genome Res 23:217–227. https://doi.org/10.1101/gr.140301.112

Kvitek DJ, Sherlock G (2011) Reciprocal sign epistasis between frequently experimentally evolved adaptive mutations causes a rugged fitness landscape. PLoS Genet 7:e1002056. https://doi.org/10.1371/journal.pgen.1002056

Laffitte M-CN, Leprohon P, Papadopoulou B, Ouellette M (2016) Plasticity of the Leishmania genome leading to gene copy number variations and drug resistance. F1000Res 5:2350. https://doi.org/10.12688/f1000research.9218.1

Lengauer C, Kinzler KW, Vogelstein B (1998) Genetic instabilities in human cancers. Nature 396:643–649. https://doi.org/10.1038/25292

Linder RA, Greco JP, Seidl F, et al (2017) The Stress-Inducible Peroxidase TSA2 Underlies a Conditionally Beneficial Chromosomal Duplication in Saccharomyces cerevisiae. G3 7:3177–3184. https://doi.org/10.1534/g3.117.300069

Oltmann J, Heselmeyer-Haddad K, Hernandez LS, et al (2018) Aneuploidy, TP53 mutation, and amplification of MYC correlate with increased intratumor heterogeneity and poor prognosis of breast cancer patients. Genes Chromosomes Cancer 57:165–175. https://doi.org/10.1002/gcc.22515

Pavelka N, Rancati G, Zhu J, et al (2010) Aneuploidy confers quantitative proteome changes and phenotypic variation in budding yeast. Nature 468:321–325. https://doi.org/10.1038/nature09529

Payen C, Di Rienzi SC, Ong GT, et al (2014) The dynamics of diverse segmental amplifications in populations of Saccharomyces cerevisiae adapting to strong selection. G3 4:399–409. https://doi.org/10.1534/g3.113.009365

Payen C, Sunshine AB, Ong GT, et al (2016) High-Throughput Identification of Adaptive Mutations in Experimentally Evolved Yeast Populations. PLoS Genet 12:e1006339. https://doi.org/10.1371/journal.pgen.1006339

Peter J, De Chiara M, Friedrich A, et al (2018) Genome evolution across 1,011 Saccharomyces cerevisiae isolates. Nature 556:339–344. https://doi.org/10.1038/s41586-018-0030-5

Potapova T, Gorbsky GJ (2017) The Consequences of Chromosome Segregation Errors in Mitosis and Meiosis. Biology (Basel) 6:12. https://doi.org/10.3390/biology6010012

Puddu F, Herzog M, Selivanova A, et al (2019) Genome architecture and stability in the Saccharomyces cerevisiae knockout collection. Nature 573:416–420. https://doi.org/10.1038/s41586-019-1549-9

Rutkowski TP, Schroeder JP, Gafford GM, et al (2017) Unraveling the genetic architecture of copy number variants associated with schizophrenia and other neuropsychiatric disorders. J Neurosci Res 95:1144–1160. https://doi.org/10.1002/jnr.23970

Scopel EFC, Hose J, Bensasson D, Gasch AP (2021) Genetic variation in aneuploidy prevalence and tolerance across Saccharomyces cerevisiae lineages. Genetics 217:iyab015. https://doi.org/10.1093/genetics/iyab015

Selmecki A, Bergmann S, Berman J (2005) Comparative genome hybridization reveals widespread aneuploidy in Candida albicans laboratory strains. Mol Microbiol 55:1553–1565. https://doi.org/10.1111/j.1365-2958.2005.04492.x

Selmecki A, Forche A, Berman J (2006) Aneuploidy and isochromosome formation in drug-resistant Candida albicans. Science 313:367–370. https://doi.org/10.1126/science.1128242

Selmecki A, Gerami-Nejad M, Paulson C, et al (2008) An isochromosome confers drug resistance in vivo by amplification of two genes, ERG11 and TAC1. Mol Microbiol 68:624–641. https://doi.org/10.1111/j.1365-2958.2008.06176.x

Sotillo R, Hernando E, Díaz-Rodríguez E, et al (2007) Mad2 overexpression promotes aneuploidy and tumorigenesis in mice. Cancer Cell 11:9–23. https://doi.org/10.1016/j.ccr.2006.10.019

Stopsack KH, Whittaker CA, Gerke TA, et al (2019) Aneuploidy drives lethal progression in prostate cancer. Proc Natl Acad Sci U S A 116:11390–11395. https://doi.org/10.1073/pnas.1902645116

Sunshine AB, Payen C, Ong GT, et al (2015) The fitness consequences of aneuploidy are driven by condition-dependent gene effects. PLoS Biol 13:e1002155. https://doi.org/10.1371/journal.pbio.1002155

Terhorst A, Sandikci A, Keller A, et al (2020) The environmental stress response causes ribosome loss in aneuploid yeast cells. Proc Natl Acad Sci U S A 117:17031–17040. https://doi.org/10.1073/pnas.2005648117

Tibshirani R, Saunders M, Rosset S, et al (2005) Sparsity and Smoothness Via the Fused Lasso. Journal of the Royal Statistical Society Series B: Statistical Methodology 67:91–108. https://doi.org/10.1111/j.1467-9868.2005.00490.x

Tong AH, Evangelista M, Parsons AB, et al (2001) Systematic genetic analysis with ordered arrays of yeast deletion mutants. Science 294:2364–2368. https://doi.org/10.1126/science.1065810

Torres EM, Dephoure N, Panneerselvam A, et al (2010) Identification of aneuploidy-tolerating mutations. Cell 143:71–83. https://doi.org/10.1016/j.cell.2010.08.038

Torres EM, Sokolsky T, Tucker CM, et al (2007) Effects of aneuploidy on cellular physiology and cell division in haploid yeast. Science 317:916–924. https://doi.org/10.1126/science.1142210

Usher CL, McCarroll SA (2015) Complex and multi-allelic copy number variation in human disease. Brief Funct Genomics 14:329–338. https://doi.org/10.1093/bfgp/elv028

Weaver BAA, Cleveland DW (2006) Does aneuploidy cause cancer? Curr Opin Cell Biol 18:658–667. https://doi.org/10.1016/j.ceb.2006.10.002

Williams BR, Prabhu VR, Hunter KE, et al (2008) Aneuploidy affects proliferation and spontaneous immortalization in mammalian cells. Science 322:703–709. https://doi.org/10.1126/science.1160058

Yang F, Todd RT, Selmecki A, et al (2021) The fitness costs and benefits of trisomy of each Candida albicans chromosome. Genetics 218:iyab056. https://doi.org/10.1093/genetics/iyab056

Yurov YB, Vorsanova SG, Demidova IA, et al (2018) Mosaic Brain Aneuploidy in Mental Illnesses: An Association of Low-level Post-zygotic Aneuploidy with Schizophrenia and Comorbid Psychiatric Disorders. Curr Genomics 19:163–172. https://doi.org/10.2174/1389202918666170717154340

