## Supplemental Figure 1 for "Condition-dependent fitness effects of large synthetic chromosome amplifications"

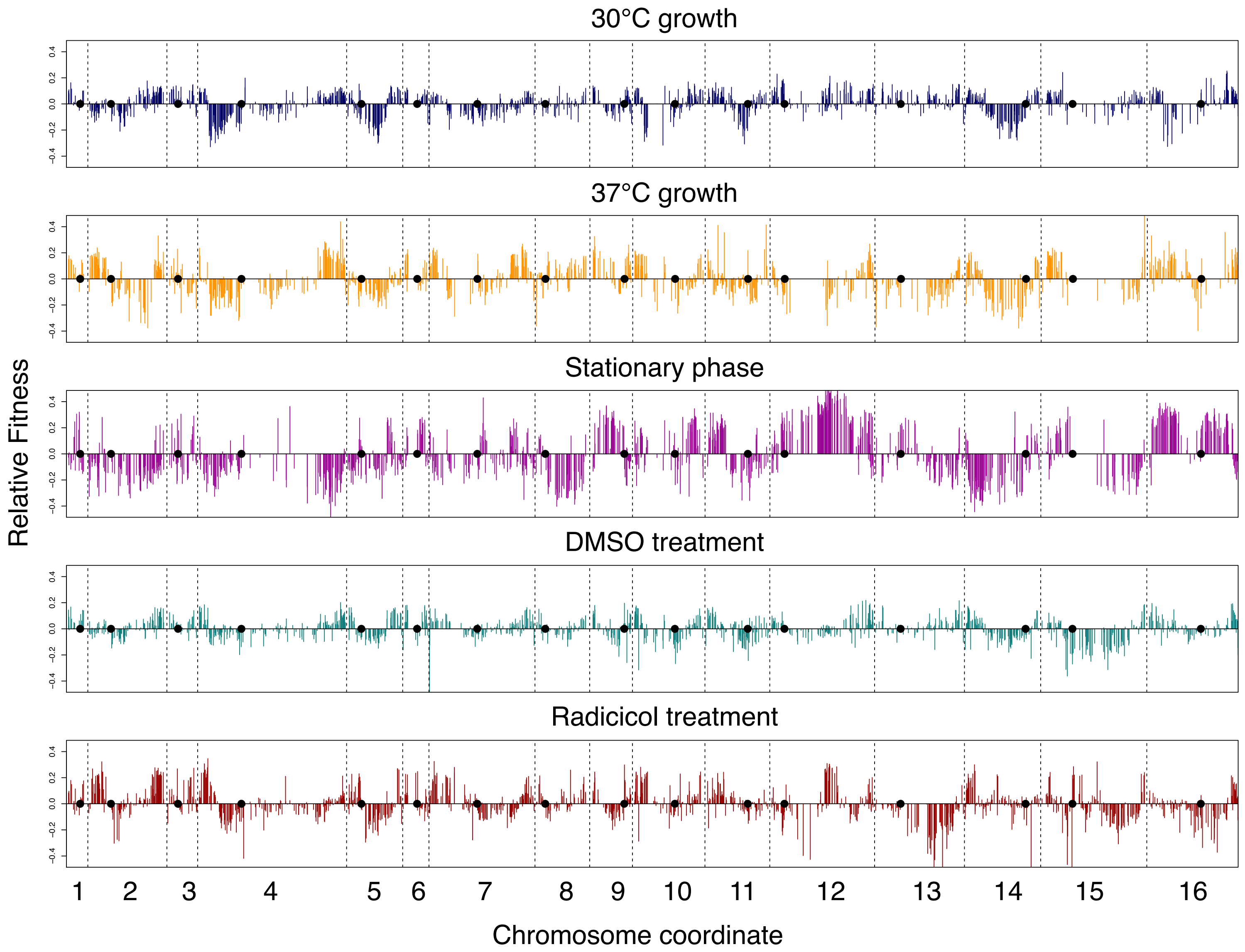


**Supplementary Figure 1: Fitness of all Tamp strains across the entire genome for each experimental condition.** Each Tamp strain is represented by a bar at the site that the Tamp initiates and extends to the telomere on the same arm. Chromosomes are delimited by dashed lines. Centromeres are represented by black dots.
