## Supplemental Figure 2 for "Condition-dependent fitness effects of large synthetic chromosome amplifications"

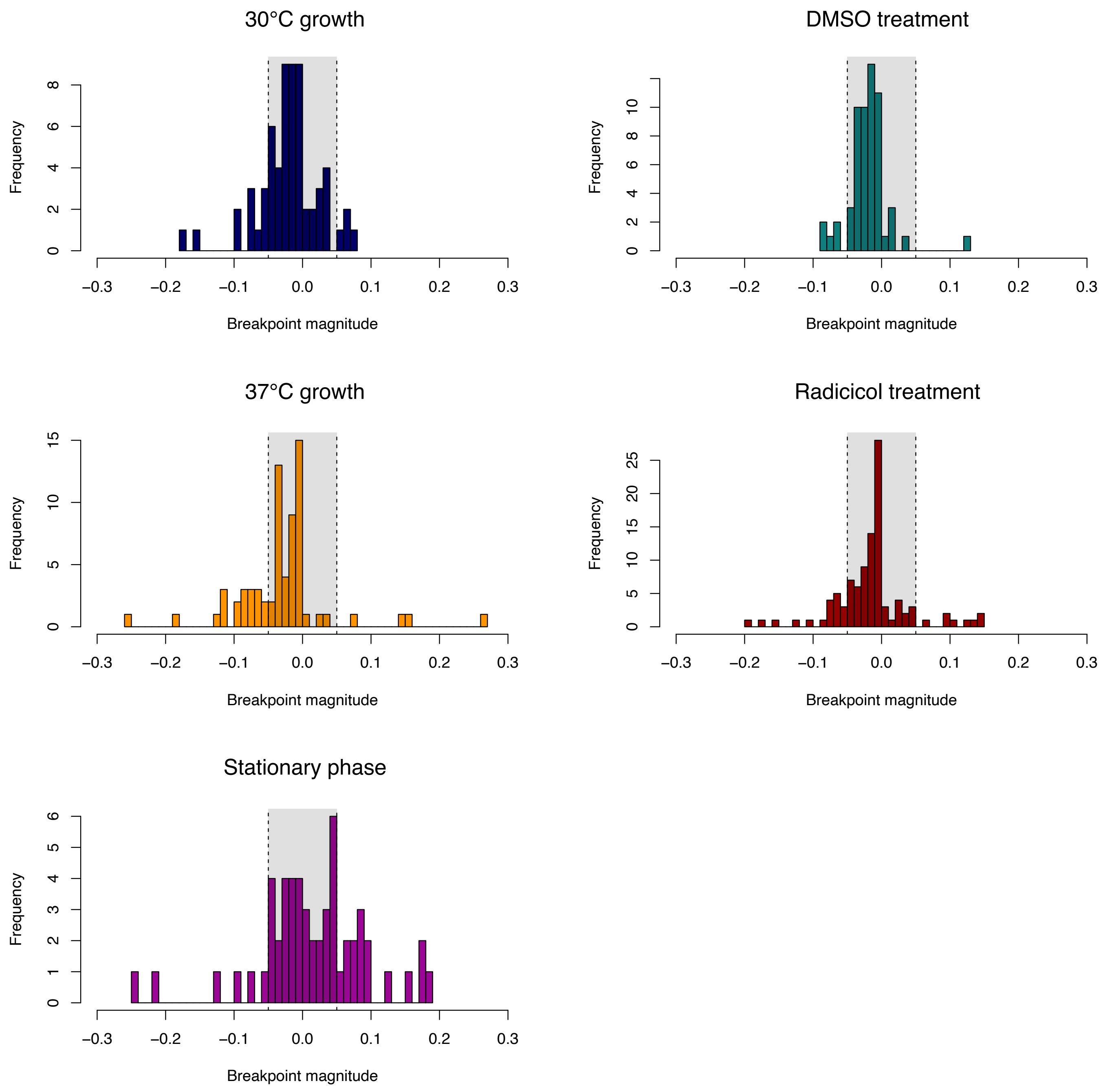


**Supplementary Figure 2: Breakpoint histograms across all conditions tested.** Breakpoint magnitude is calculated as the magnitude of the step points of the piecewise constant model. Upsteps, regions that improve fitness when amplified have a positive magnitude and downsteps and negative magnitude. We narrowed our candidate list of breakpoints as those outside our +/- 0.05 cutoff (dotted lines).
