## Supplemental Figure 3 for "Condition-dependent fitness effects of large synthetic chromosome amplifications"

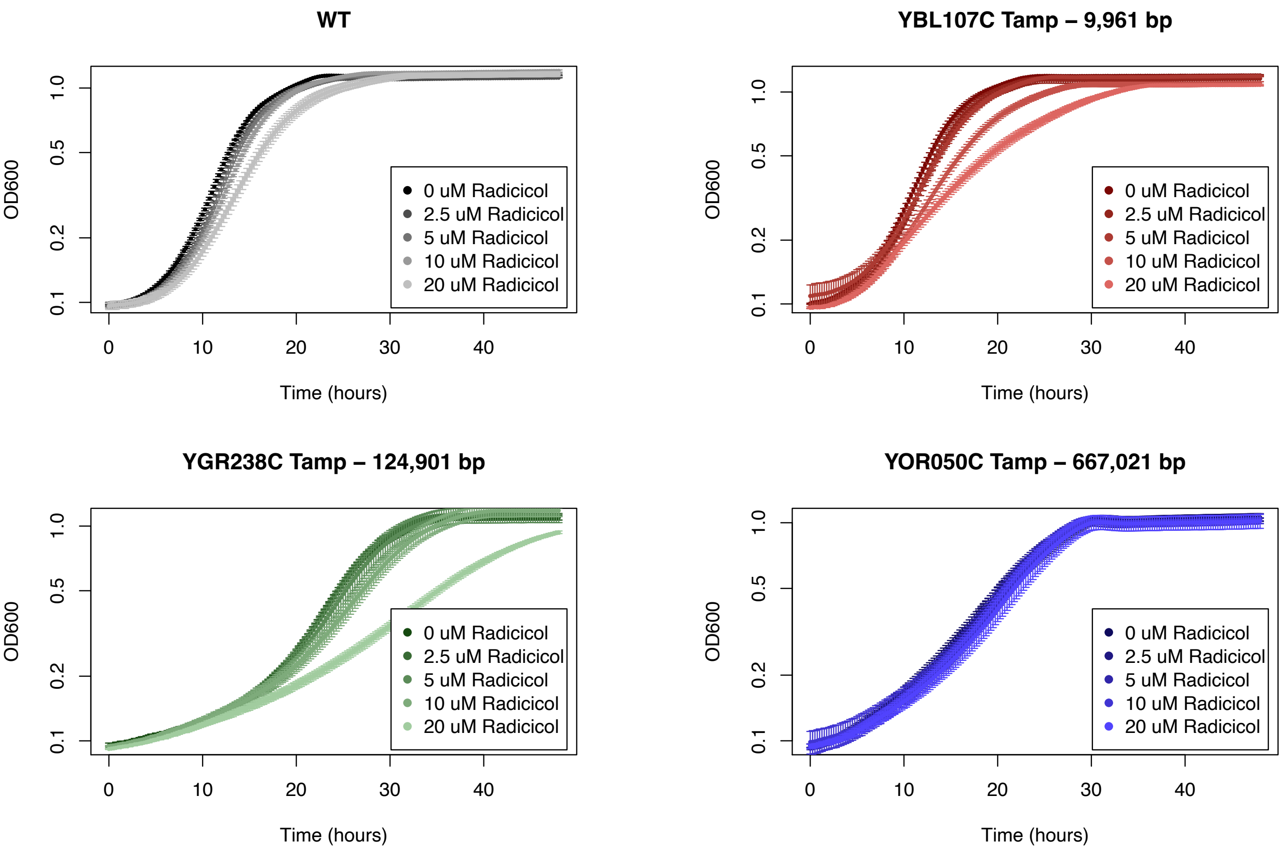


**Supplemental Figure 3: Effect of radicicol treatment on wild type and Tamp strain growth.** Strains were grown in triplicate in a BioTek Synergy H1 Multi-Mode plate reader. Despite YOR050C being the longest Tamp that should be the most sensitive, strains disomic for chromosome 11 do not show sensitivity to Hsp90 inhibitor.
